# ‘Ripple effects’ of urban environmental characteristics on cognitive performances in Eurasian red squirrels

**DOI:** 10.1101/2022.04.20.488863

**Authors:** Pizza Ka Yee Chow, Kenta Uchida, Itsuro Koizumi

**Affiliations:** Department of Psychology, University of Chester, UK; Ecology and Genetic Research Unit, University of Oulu, Finland; Graduate School of Environmental Science, Hokkaido University, Japan; Graduate School of Agricultural and Life Sciences, University of Tokyo, Japan

**Keywords:** behavioural flexibility, cognitive ecology, domino effect, human-induced rapid environmental change, urbanisation, urban ecology, urban wildlife, wildlife behaviour

## Abstract

1. Urban areas are expanding exponentially, leading more wildlife species to reside and settle in this environment. Urban environmental characteristics, such as human disturbance or green coverage, have been shown to affect some cognitive abilities such as innovative problem-solving performance of wildlife species. However, an untested hypothesis is that due to the shared underlying cognitive mechanisms, these affected performances may induce a ‘ripple’ effect, and continue to affect other related cognitive processes (the ripple effect hypothesis).
2. We tested this hypothesis by targeting two cognitive abilities, generalisation and memory, that overlap the cognitive mechanisms (learning and memory) of the original problem solving task in urban Eurasian red squirrels. These squirrels reside in 11 urban areas where they had previously repeatedly solved the original task (the innovators), and that their solving performance in the original task was affected by the selected urban environmental characteristics. We presented two established food-extraction tasks to the innovators to measure their performance in applying the learned successful solutions when solving a similar but novel problem (i.e., generalisation process) and recalling the learned solution of the original problem when solving the same task after an extended period of time (i.e., memory).
3. Our results provide more detailed information to refine the hypothesis; the initial effects of urban environmental characteristics on the performance of the original task affect performance at individual level but not at population level. These affected performance includes individuals’ generalisation solving latency across successes as well as their first solving latency in the memory task.
4. Urban environmental characteristics affect solving performance at both population and individual levels. Some environmental characteristics such as direct and indirect human disturbance affect the success of solving the generalisation task and the memory task at site level whereas other environmental characteristics such as green coverage affect the individuals’ solving latency in both tasks.
5. Overall, our results support the ripple effect hypothesis, indicating that urban environmental characteristics have a more global impact on shaping cognitive performance than previously has shown, and thus provide a better understanding of the mechanism that supports wildlife in adapting to urban environments.

## Introduction

Urban areas are growing exponentially, leading more wildlife species to reside in urban environments as their alternative habitats. Mounting evidence has shown that urban environments have changed wildlife’s fitness-related traits, such as in physiology (Patankar et al., 2021), morphology (Biard et al., 2017; Evans et al., 2009), and behaviour (Lowry et al., 2013; Ritzel & Gallo, 2020; Santini et al., 2019; Sih, 2013; Sih et al., 2011; Sol et al., 2013). Despite these investigations, it is largely unclear how urban environments shape wildlife cognition (Griffin et al., 2017; Lee & Thornton, 2021) which can be considered as the process of an individual acquiring, storing, utilising and reacting to their environment (Shettleworth, 2010). The fact that cognition manifests via behaviour suggests that these two mechanisms have a tight relationship (Shettleworth, 2010), and that cognition, like behaviour, is a crucial fitness-related trait that facilitates wildlife adapting to urban environments (Sol et al., 2008, 2013). Consequently, investigations in this area are necessary to highlight the role of urban environments in shaping cognition, which will yield a better understanding about the mechanisms supporting wildlife species to thrive or decline in urban environments (Sol et al., 2008).

Investigations in urban wildlife cognition is still in infancy (Lee & Thornton, 2021), but studies have suggested that urban wildlife shows enhanced ability to innovate or are more successful to use alternative solutions to obtain food from novel apparatus (Audet et al., 2016; Ducatez et al., 2015; Mazza & Guenther, 2021; Overington et al., 2009; Papp et al., 2015; Preiszner et al., 2017). A few recent studies have further shown that innovative problem-solving ability is directly affected by urban environmental characteristics such as human disturbance. For example, when human disturbance is measured as the presence of a human, it decreases the innovation success rate of wild-caught, captive urban House finches, *Haemorhous mexicanus* (Cook et al., 2017). With this in mind, the understanding of how urban environmental characteristics affect innovative problem-solving performance is far from complete. The primary reason is because a cognitive ability and its processes often overlap with other cognitive abilities (Wang & Chiew, 2010). The relatedness between cognitive abilities indicates that they share an underlying mechanism. Using problem-solving ability as an example, learned a solution to solve a problem can be related to the ability to apply the learned information to solve a similar but novel problem (i.e., generalisation) or the ability to recall the relevant or learned information in the face of the same problem (i.e., memory). Thus, learning and memory are the mechanisms that share among these cognitive abilities. The overlapped cognitive abilities suggest an obvious, but untested, hypothesis that if environmental characteristics have affected one ability (Fig. 1 path i), then this affected ability could initiate a ‘ripple effect’ (cascading effect) whereby the affected performance would have an impact on other related abilities (Fig. 1 from path i to ii, or from path i to iii) (hereafter, the ripple effect hypothesis). Note that this ripple effect is alongside the direct effects that urban environmental characteristics have on the related processes (Fig. 1 path iv and v).

**Figure 1.**
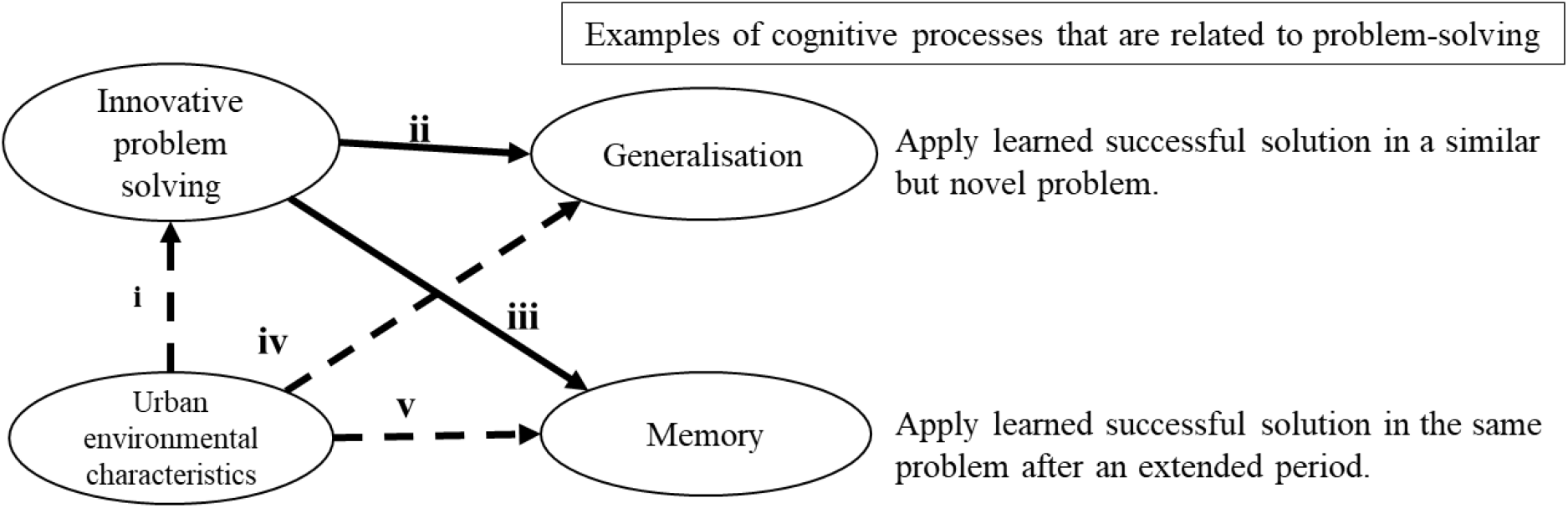
The ripple hypothesis proposes that if the urban environmental characteristics have affected one cognitive ability (indicate as innovative problem solving, path i), then the affected performance would continue to influence other related cognitive abilities, demonstrated as generalisation (path i-ii) and memory (path i-iii) due to the shared underlying cognitive mechanisms. Such a ‘ripple’ effect is additional to the direct effects that urban environmental characteristics would induce on these related processes (iv and v). See Figure 2 for a detailed model explanation.

In this study, our major goal was to test the ripple effect hypothesis (Fig. 2). To do so, we carried out a field experiment based on our previous study that has examined the effects of four urban environmental characteristics (direct human disturbance, indirect human disturbance, squirrel population size and area of green coverage) on the innovative problem-solving performance of a successful urban-dwelling species, the Eurasian red squirrels (*Sciurus vulgaris*) (Chow et al., 2021). We show that urban environmental characteristics have affected innovative problem-solving performance at population (i.e., the proportion of success) and individual level (i.e., squirrels’ solving latency on their first success or ‘first original latency’, and solving latency over time or ‘original latency across successes’) (Fig. 2 path a). Following our proposed hypothesis, any of these affected performances could be the beginning of the ripple effect and affect the performance of other related abilities such as generalisation and memory (Fig. 2, path a-b, path a-c, path a-b-d), Simultaneously, the urban environmental characteristics would also exert their direct effects on the related cognitive processes (Fig. 2 path e and path f). By relating these initial affected performances to the corresponding performances of other related abilities (see analyses for details), we will gain a deeper understanding about whether the initial effects of urban environmental characteristics have a more global impact on cognition.

**Figure 2.**
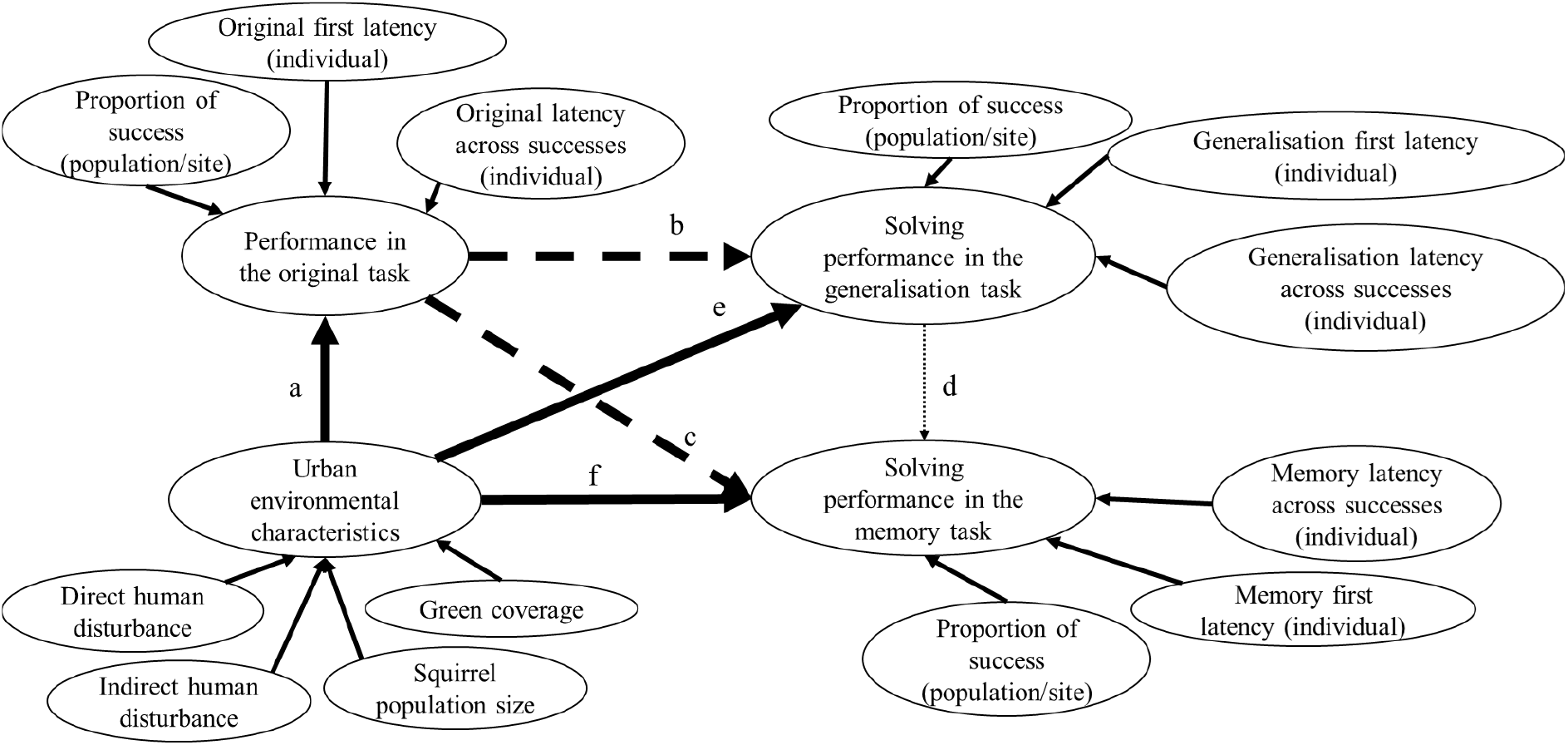
Proposed models of the ripple effect hypothesis. Path a includes the initial direct effects of urban environmental characteristics on the solving performance in the original task (assess innovative problem solving). The urban environmental characteristics were the four previously identified characteristics that include direct human disturbance, indirect human disturbance, squirrels population size and area of green coverage (m2). The effects of these characteristics on the solving performance in the original task include solving success at the population level (proportion of success), solving latency on the first success at individual level (original first latency) or solving latency across successes at individual level (original latency across successes). The initial effects become the beginning of the ripple effect that will influence the performances in the generalisation task (path a-b) or the memory task (patha-c) due to shared underlying cognitive mechanisms such as learning and memory. If the generalisation task and the memory task also share the same underlying mechanism, then the ripple effect will carry on from the generalisation task to the memory task (path a-b-d). Aside from the ripple effect, urban environmental characteristics will also affect the performance in the generalisation task (path e) and the memory task (path f). Performance in the generalisation and memory tasks can be measured at population or individual level. These measures include proportion of success at population (site) level, generalisation or memory first latency at individual level and (generalisation or memory) latency across successes at individual level.

The designs to test this hypothesis require tasks that share an underlying cognitive mechanism to the original task, which is learning and memory. The performance of generalisation and memory can be assessed using two established food-extraction tasks whereby these tasks required the squirrels that have solved the original task more than once (hereafter, the innovators) to apply, or recall, the learned successful solutions of the original task (Chow et al., 2017) (see Materials and Methods for details). Specifically, the generalisation performance is assessed by using a similar but novel apparatus that requires the innovators to apply the learned successful solutions of the original task (Fig. 3A). The memory performance is assessed by using the original task and requires the innovators to recall the learned successful solutions of the original task when solving the same task after an extended period (Fig. 3B). In both tasks, solving performances are quantified in three measures: the proportion of success at population (site) level, solving latency on the first success as well as solving latency across successes at individual level. Each of these measures can then be related to the corresponding affected performance in the original task. If the ripple effect hypothesis is true, then the initial solving performances in the original task that have been affected by the urban environmental characteristics would have an impact on the performances of the generalisation task (Fig. 2 path a-b) and the memory task (Fig. 2 path a-c).

**Figure 3.**
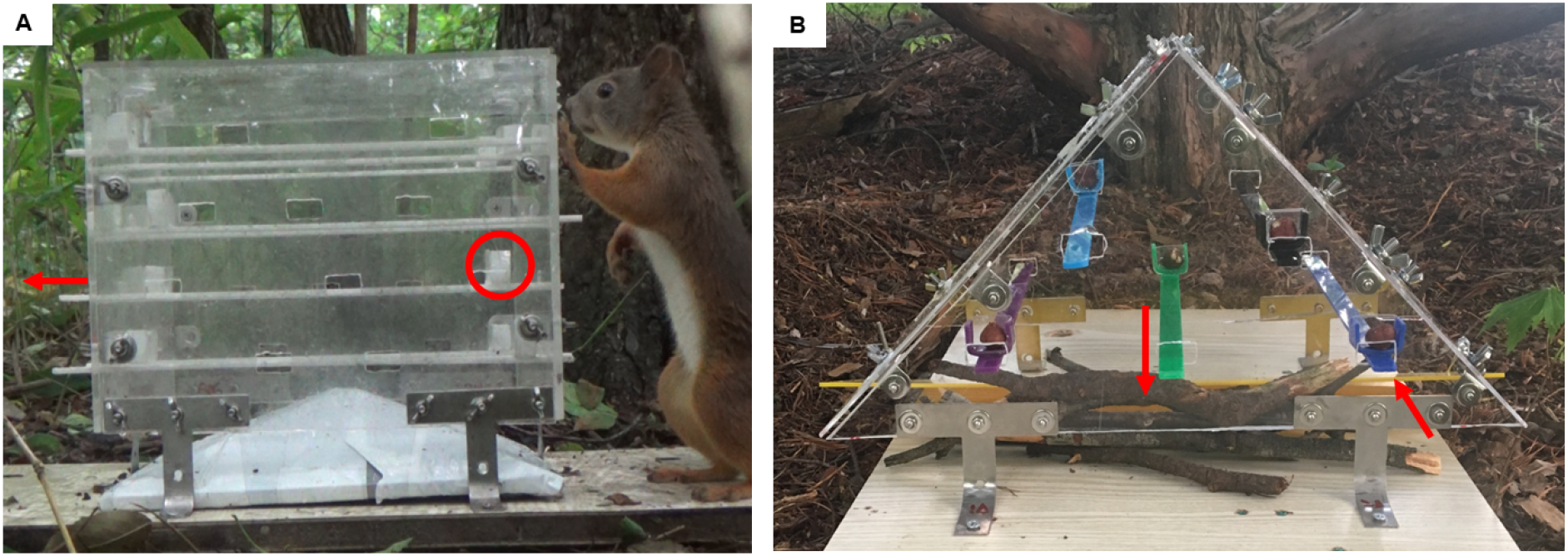
Two established food-extraction tasks were given to urban wild red squirrels. Both tasks contained the same successful solving solution that required squirrels to either push a lever if they are close to a nut container or pull a lever if they are far from the nut container. **A)** the original task that was used to measure innovative problem solving performance in urban squirrels (Chow et al., 2021). This task was presented to the innovators (the squirrels that had solved the task repeatedly) 21 days after the generalisation task to measure memory of the successful solution of the original task. **B)** the generalisation task that was used to measure the ability to apply the learned successful solutions of the original task in this similar but different task.

The effects of urban environmental characteristics on other related cognitive processes have not been tested. Here, we examined the effect of the previously identified urban environmental characteristics (i.e., direct human disturbance, indirect human disturbance, squirrel population size and green coverage) on the performances of the generalisation task and the memory task. Due to the shared underlying mechanisms, we expected some of these environmental characteristics would affect the performance of either task (Fig. 2 path e and f), both at population (site) and individual level.

## Materials and Methods

Previously, Chow and colleagues (2021) examined the innovative problem-solving performance of 71 red squirrels in 11 urban areas at Obihiro city, Hokkaido, Japan (Table S1). These areas were located at different parts of the city (> 800 m between sites to avoid pseudo-replication) and varied in their environmental characteristics. Of these squirrels, 38 of them (53.5%) were innovators that had solved the task repeatedly, which allowed us to test the ripple effect hypothesis with them. Between May 2018 and January 2019, we used the same location that was used to present the original task to test our hypothesis. Each of these locations was selected for safety purposes; the locations were distant from major roads and close to trees (to avoid road kill and predation risk, see detail in Table S1). We presented the tasks daily during the innovators’ most active period (from dawn to noon) regardless of weather conditions. We checked (i.e., refilled the apparatus and carried out randomisation procedures) the site 3-4 times per day (45 mins-1.5 hours between checks). All squirrels were identified by an established method (Chow et al., 2018) that involved frame-by-frame analysis of each squirrel’s physical characteristics from video footage as well as mark-recaptured and mark-resight methods (see Note S1). This study was approved by Hokkaido University (ethics number: 606) and Obihiro University.

### (a) Urban environmental characteristics

Four previously identified environmental characteristics and measurements were adopted for this study (Chow et al., 2021) (see Note S2 for more detailed information regarding each measurement): 1) direct human disturbance (mean number of humans in a site per day that was the average of 4-5 times walking in a site, which included before or after checking/refilling the apparatus across all observation days); 2) indirect human disturbance (number of buildings including houses and stores 50 m around a site); 3) area of green coverage in metres (total area covered by trees in a site (m^2^) on Google satellite; and 4) squirrel population density (number of squirrels in each site divided by the site size (m^2^) from regular field surveys, trapping records, and video footage).

### (b) Food-extraction tasks

The original task had a cube-shaped top and a pyramid-shaped bottom (Fig. 3A). The top was secured by six steel legs, creating a 3.5 cm gap between the top and the bottom for the innovators to receive a reward (hazelnut) upon successfully solving the task. Each side of the top had ten horizontally, but not vertically, aligned holes. These holes were roughly aligned to the holes on the opposite side. We inserted ten (white) levers horizontally across the box through the holes, leaving 2.5 cm of both lever-ends outside the holes so that squirrels can manipulate the lever ends. Each lever had a nut container 2.5 cm away from one of its ends, which can be positioned inside the box. To solve this task, an innovator needed to either push a lever end if it was close to a nut container, or pull the lever end if it was far from the nut container (video S1: original task). With repeated successes of extracting nuts from the apparatus, the innovators become more efficient in deploying the pull-or-push solutions (Chow et al., 2021).

To assess the generalisation process, we presented to the innovators a similar but novel food-extraction apparatus (hereafter, the generalisation task) (Chow et al., 2017) (Fig. 3B; video 2: generalisation task). The main feature of this task was that it had the same successful solving solutions as the original task (i.e., we used levers to facilitate the application of the learned successful solutions), but the apparatus had a different shape and colour (triangle front: 35 × 19 × 18 cm; length x width x height, rectangular side: 25 × 20 cm) to maximise the difference between this task and the original task. This apparatus had a transparent top and a rectangular plexiglas base. The top was secured by four steel legs, creating a 3 cm gap for the squirrels to retrieve a nut on successful solving. The front and the back of the top had five holes that were horizontally, but not vertically, aligned with each other. Five levers could be inserted into the apparatus across the holes. Each lever had a nut container located 1.5 cm away from one lever-end and positioned just inside the box. The levers and their facing direction in the box were randomly presented in each check.

We examined the innovators’ memory for the learned successful solutions of the original task 21 days after the generalisation task. To do so, we presented the original apparatus (Fig. 3A) to the innovators for one field day. Between the generalisation task and the memory task, we kept the innovators visiting the location by carrying out other behavioural assays that did not involve any similar solutions for solving the generalisation or the memory task. As with the original task, we randomised the facing direction of the box and the levers as well as randomly choosing which levers contained a nut in the memory task (see Note S3 for detailed experimental protocol).

### (c) Behavioural measurements

#### Affected solving performance and other relevant variables in the original task

The following solving performances in the original task were affected by the selected urban environmental characteristics, and each of these affected performances could be the start of the ripple effect: 1) success rate at the population (site) level; 2) the first original latency at the individual level; and 3) original latency across successes at individual level. Due to the design of the generalisation and the memory tasks containing the same solving solutions, we also recorded additional variables that may affect the performance in either task. For the generalisation task, these variables included the total number of successes each innovator obtained in the original task and their last original solving latency. For the memory task, related variables included the number of successes that each innovator obtained both in the original task and the generalisation task, the last original solving latency, the first generalisation solving latency and the last generalisation solving latency.

#### Solving performance in the generalisation and memory task

When an innovator attempted to solve a task (indicated as using their paw, nose or incisors to manipulate a lever), we recorded the time that the attempt started until the innovator stopped doing so (i.e., stopped manipulating the level). This record included two information: the solving outcome (whether the innovator successfully solved a task, succeeded or failed) and solving duration per attempt regardless of whether that attempt led to a successful or failed solving.

For each innovator, we recorded whether it solved the task on its first visit (recorded as an innovator first appeared on a video until it left the apparatus for more than 2 minutes) or on its subsequent visits (recorded as the innovator reappeared on the videos). We considered each innovator as a ‘problem solver’ if it solved the generalisation task (presented in a novel apparatus) or the memory task (presented in the original apparatus) more than once. Others were classified as ‘non-problem solvers’ if the innovators only solved the task once but could not repeat the success throughout the task presentation. We calculated the proportion of success at site level (i.e., dividing the number of problem solvers successfully solving a task by the number of innovators participating in the task for each site). For each problem solver, we calculated the solving latency to each success (i.e., summing all solving latencies of unsuccessful attempts until a success occurred) in order to obtain the solving latency for the first success and each subsequent success. We referred the latency in the original task to ‘original latency’, ‘generalisation latency’ for the generalisation task and ‘memory latency’ for the memory task.

We additionally recorded the solution that an innovator used to solve the task the first time; this gave information on whether the innovators had applied the learned successful solution of the original task when they were solving the generalisation task, or when they re-encountered the original task after an extended time period. For example, if an innovator had used the same successful solving solution to solve the generalisation task and the memory task, then this would indicate their successful application or recall of the learned information. In this case, their first (generalisation or memory) solving latency in each task would be significantly different from their first original latency (i.e., when the innovators had not learned the successful solution in the original task) but comparable to their last original latency (when the innovators had learned the solution). However, if the innovators considered the generalisation task as a new task, or that they had forgotten the solution of the original task, then their solution to solve the generalisation task and the memory task would be different from the very last solution that they used to solve the original task. Their first solving latency in each task would also be comparable to the first original latency as well as significantly different from their last original latency.

### (d) Data analysis

We analysed data using R (version 3.5.2) (Ripley, 2001) and SPSS (version 25). We report the original values of mean (M) ± standard errors (S.E). A two-tailed test with α ≤ 0.05 is considered as significant. As our aim was on testing the hypothesis, multicollinearity between variables was less of a concern. We standardised the variables for model convergence purposes. Model fit was shown using marginal R^2^ (R^2^m) which higher marginal R^2^ indicated the selected variables in a model had higher variance. For each task, beta regression in the package ‘betareg’ (Cribari-Neto & Zeileis, 2010) was used to model the proportion of success at population (site) level whereas Generalized Linear Mixed Models (GLMM) with gamma log link distribution in the package ‘glmmTMB’ (Magnusson et al., 2017) was used to accommodate positively skewed continuous response variables (i.e., first solving latency and solving latency across successes).

We first examined general task performance for each task. We checked whether each problem solver used its last solving solution in the original task to solve the generalisation and memory tasks. For the generalisation task, we compared the problem solvers’ first generalisation latency with their first and last original latency as a reflection on whether they could quickly apply the learned solutions to solve the generalisation task. As most of the problem solvers had solved the task 10 times in the generalisation task, we also examined generalisation latency across 10 successes to assess their learning performance in this task. For the memory task, we compared each problem solver’s first memory latency with their first and last original latency in the original task to assess their memory of the learned successful solutions of the original task. As the generalisation task also had the same solving solution as the original and the memory tasks, the solving performance in the generalisation task may affect the solving latency in the memory task. Accordingly, we also compared the memory latency with the first and last generalisation latency. Most of the problem solvers in the memory tasks obtained 5 successes, and we examined recall latency across the 5 successes.

We then examined the ripple effect hypothesis. To do this, we ran two models that included the direct effects of urban environmental characteristics on two solving performances of the original problem (Path a in Fig. 2). Each model included four environmental characteristics as fixed factors (i.e., direct human disturbance, indirect human disturbance, green coverage (m^2^), and squirrel population size), and a solving performance as response variable (i.e., the proportion of success at the population (site) level and the first original solving latency at individual level). In the first original latency model, we also included site and individual identity as random variables.

For the generalisation task, we ran three models to assess whether the initial affected performance (along with the four urban environmental characteristics) influenced each of the generalisation performance (success rate at site level, first generalisation latency, and generalisation latency across successes) (Path b & e in Fig. 2). When modelling the proportion of success for the generalisation task, the fixed variables included the four urban environmental characteristics and the previous proportion of success at population (site) level in the original task. When modelling the first generalisation latency and generalisation latency across successes, the first and the last original latency, the number of successes that each problem solver obtained in the original task and the four urban environmental characteristics were included as fixed variables (See Table 1 for the list of fixed variables in each model). In these models, site and individual identity were included as random variables.

**Table 1.** Problem-solving performances of the generalisation task. All models include four urban characteristics alongside solving performance(s) or related variables of the original task as fixed factors. Response variable of each model is listed on the left side of the table. This table includes estimates (Est), standard errors (S. E.), Z values, P values, Marginal R^2^ (R^2^m). Bolded values indicate a fixed variable is P ≤ 0.05. Higher value of R^2^m indicates the variables in the model have higher variance.

|  | Response variable | Fixed variables | Est | S. E. | Z | P | R<sup>2</sup>m |
| --- | --- | --- | --- | --- | --- | --- | --- |
| a | Generalisation success | Direct human disturbance | -1.97 | 0.67 | -2.94 | **0.003** | 0.75 |
|  |  | Indirect human disturbance | -1.67 | 0.46 | -3.67 | **<0.001** |  |
|  |  | Squirrel population size | 0.19 | 0.42 | 0.45 | 0.654 |  |
|  |  | Green coverage | -0.39 | 0.34 | -1.13 | 0.259 |  |
|  |  | Solving success rate on the original task (at site level) | -1.05 | 0.61 | -1.73 | 0.084 |  |
| b | Generalisation latency on the first success | Direct human disturbance | 0.27 | 0.23 | 1.18 | 0.239 | 0.13 |
|  |  | Indirect human disturbance | 0.05 | 0.32 | 0.16 | 0.877 |  |
|  |  | Squirrel population size | 0.33 | 0.28 | 1.17 | 0.243 |  |
|  |  | Green coverage | 0.17 | 0.31 | 0.55 | 0.581 |  |
|  |  | Innovator's first solving latency in the original task | 0.18 | 0.26 | 0.70 | 0.485 |  |
|  |  | Innovator's last solving latency in the original task | -0.09 | 0.26 | -0.33 | 0.739 |  |
|  |  | Number of successes obtained in the original task | -0.04 | 0.28 | -0.14 | 0.885 |  |
| c | Generalisation latency across successes | Direct human disturbance | 0.14 | 0.09 | 1.52 | 0.128 | 0.13 |
|  |  | Indirect human disturbance | -0.05 | 0.10 | -0.52 | 0.602 |  |
|  |  | Squirrel population size | 0.06 | 0.09 | 0.72 | 0.469 |  |
|  |  | Green coverage | 0.25 | 0.09 | 2.63 | **0.008** |  |
|  |  | Innovator's first solving latency in the original task | 0.24 | 0.08 | 3.03 | **0.002** |  |
|  |  | Innovator's last solving latency in the original task | 0.06 | 0.09 | 0.70 | 0.483 |  |
|  |  | Number of successes obtained in the original task | -0.15 | 0.08 | -1.77 | 0.076 |  |
|  |  | Success number | -0.13 | 0.03 | -4.76 | **<0.001** |  |

For the memory task, we also ran three models to examine whether the initial affected performance (alongside the four urban environmental characteristics) influenced the performance in the memory task (Path c in Fig. 2). When modelling the proportion of success of the memory task, fixed variables included the previous proportion of success in the original task and the four urban environmental characteristics. When modelling the first memory latency and memory latency across successes, we not only included the first and last original latency, but also included the first and last generalisation latency as well as the total number of successes obtained in both the original task and the generalisation task in the model (see Table 2 for the list of fixed variable in each model). Site was the only random variable for all these models due to convergence issues.

**Table 2.**
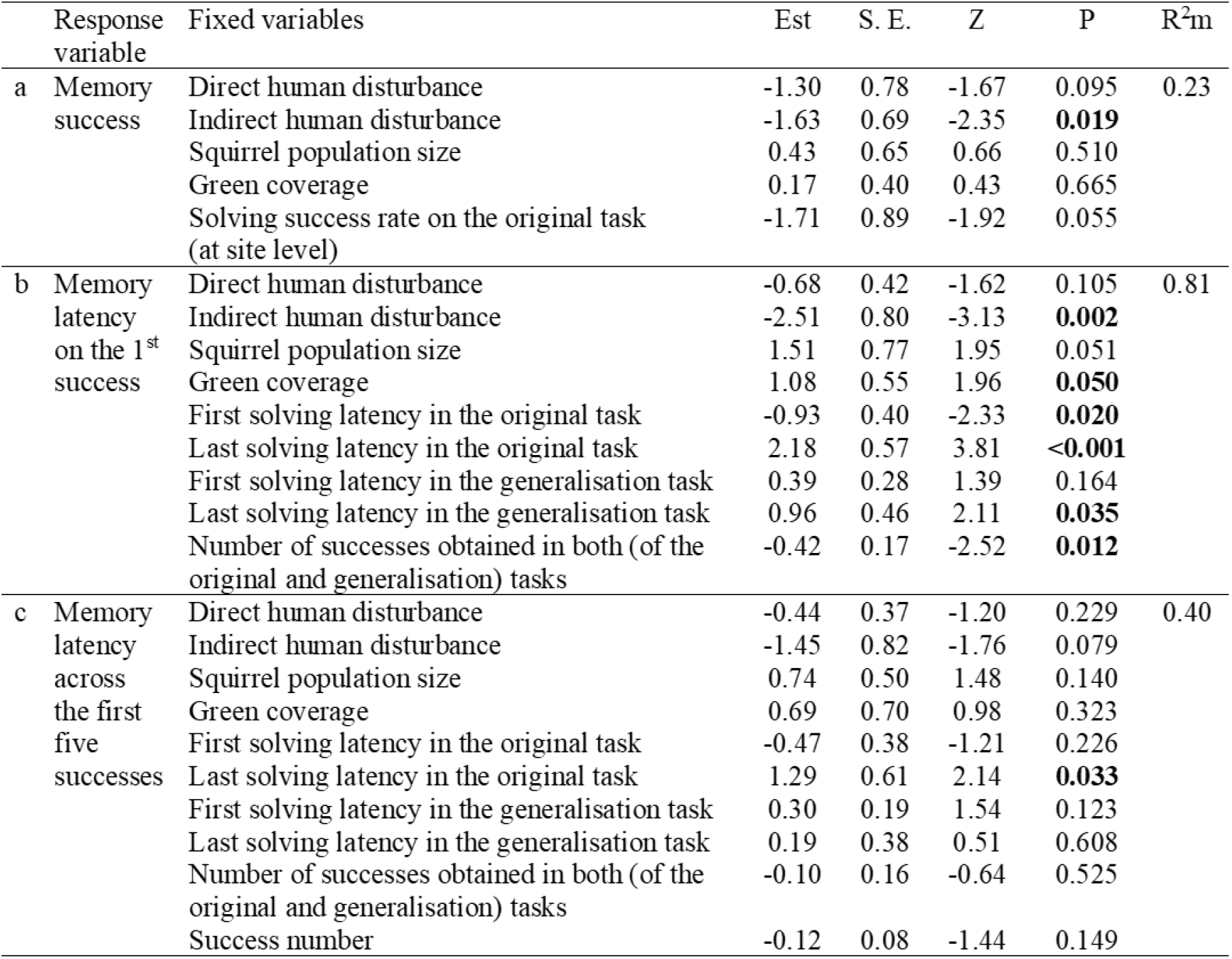
Problem-solving performances of the memory task. All models include four urban characteristics alongside solving performance(s) or related variables of the original task as fixed factors. Response variable of each model is listed on the left side of the table. This table includes estimates (Est), standard errors (S. E.), Z values, P values, Marginal R^2^ (R^2^m). Bolded values indicate a fixed variable is P ≤ 0.05. Higher value of R^2^m indicates the variables in the model have higher variance.

| Response variable | Fixed variables | Est | S. E. | Z | P | R<sup>2</sup>m |
| --- | --- | --- | --- | --- | --- | --- |
| a Memory success | Direct human disturbance | -1.30 | 0.78 | -1.67 | 0.095 | 0.23 |
|  | Indirect human disturbance | -1.63 | 0.69 | -2.35 | **0.019** |  |
|  | Squirrel population size | 0.43 | 0.65 | 0.66 | 0.510 |  |
|  | Green coverage | 0.17 | 0.40 | 0.43 | 0.665 |  |
|  | Solving success rate on the original task (at site level) | -1.71 | 0.89 | -1.92 | 0.055 |  |
| b Memory latency on the 1<sup>st</sup> success | Direct human disturbance | -0.68 | 0.42 | -1.62 | 0.105 | 0.81 |
|  | Indirect human disturbance | -2.51 | 0.80 | -3.13 | **0.002** |  |
|  | Squirrel population size | 1.51 | 0.77 | 1.95 | 0.051 |  |
|  | Green coverage | 1.08 | 0.55 | 1.96 | **0.050** |  |
|  | First solving latency in the original task | -0.93 | 0.40 | -2.33 | **0.020** |  |
|  | Last solving latency in the original task | 2.18 | 0.57 | 3.81 | **<0.001** |  |
|  | First solving latency in the generalisation task | 0.39 | 0.28 | 1.39 | 0.164 |  |
|  | Last solving latency in the generalisation task | 0.96 | 0.46 | 2.11 | **0.035** |  |
|  | Number of successes obtained in both (of the original and generalisation) tasks | -0.42 | 0.17 | -2.52 | **0.012** |  |
| c Memory latency across the first five successes | Direct human disturbance | -0.44 | 0.37 | -1.20 | 0.229 | 0.40 |
|  | Indirect human disturbance | -1.45 | 0.82 | -1.76 | 0.079 |  |
|  | Squirrel population size | 0.74 | 0.50 | 1.48 | 0.140 |  |
|  | Green coverage | 0.69 | 0.70 | 0.98 | 0.323 |  |
|  | First solving latency in the original task | -0.47 | 0.38 | -1.21 | 0.226 |  |
|  | Last solving latency in the original task | 1.29 | 0.61 | 2.14 | **0.033** |  |
|  | First solving latency in the generalisation task | 0.30 | 0.19 | 1.54 | 0.123 |  |
|  | Last solving latency in the generalisation task | 0.19 | 0.38 | 0.51 | 0.608 |  |
|  | Number of successes obtained in both (of the original and generalisation) tasks | -0.10 | 0.16 | -0.64 | 0.525 |  |
|  | Success number | -0.12 | 0.08 | -1.44 | 0.149 |  |

## Results

For the generalisation task, 28 (out of 38, 74% participation rate) innovators who had repeatedly successfully solved the original task participated in this task. 23 (82.1%) innovators from 10 sites eventually solved the task in the first or subsequent visits, and thus were the ‘problem solvers’ for this task. In this task, the proportion of success at site level (P = 0.084) was not affected by a site’s previous success rate but by urban environmental characteristics that included mean number of humans in a site per day (beta regression: *Z* = −2.94, P = 0.003) and the number of buildings around a site (*Z* = −3.67, P < 0.001) (Table 1a). Increased (direct and indirect) human disturbance decreased success rate at site level. When the problem solvers solved the generalisation task the first time, their first generalisation latency was significantly longer than their last original latency (GLMM: *Z* = −2.29, P = 0.022), and comparable to the first original latency (*Z* = 1.86, P = 0.062, Fig. 4A). This first generalisation latency was not affected by any variables that were included in the model (Table 1b).

**Figure 4.**
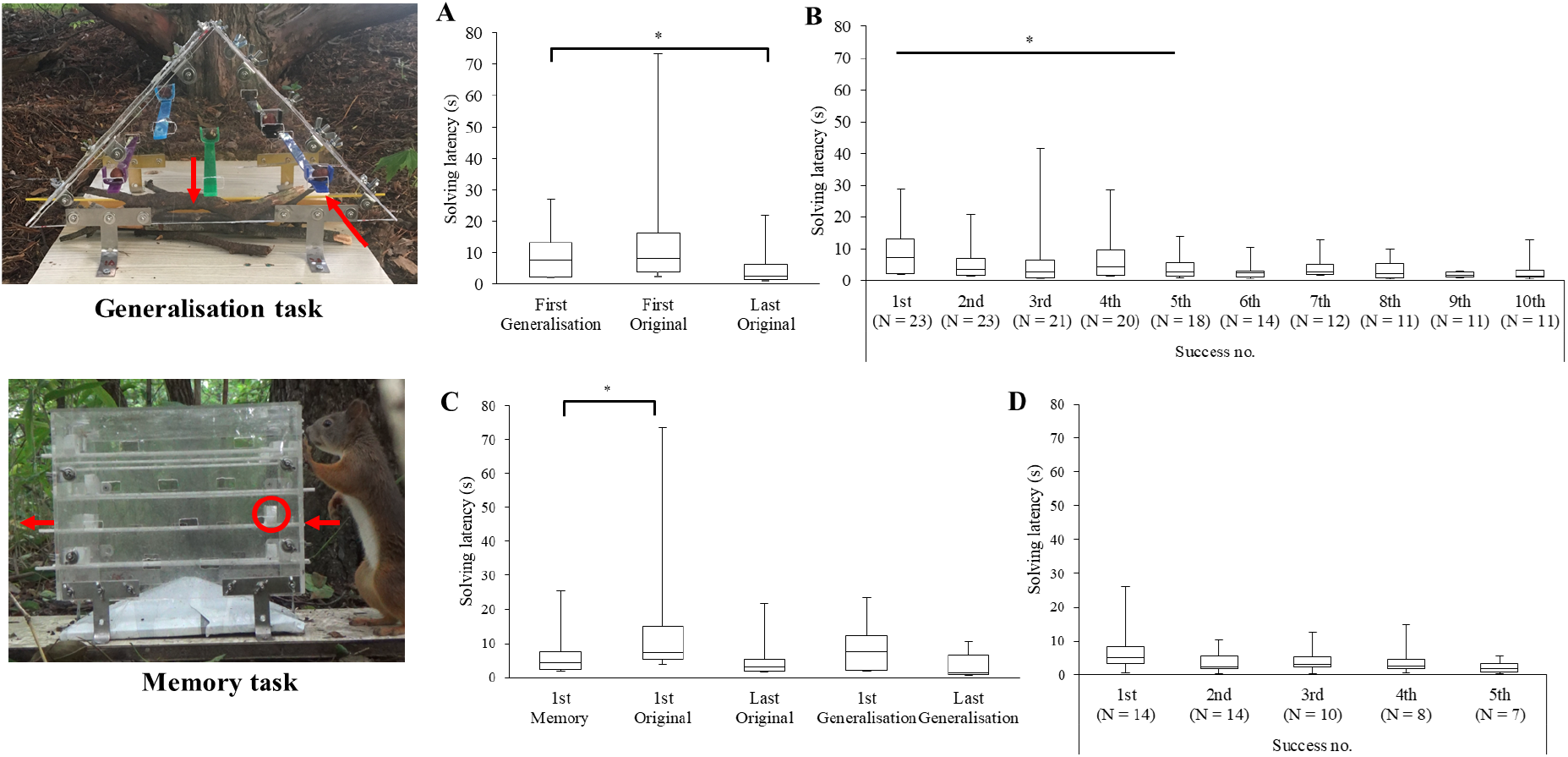
Solving latency in the generalisation task and the memory task. **A)** the first generalisation latency in seconds compared with the first and the very last original latency (seconds) (N = 23). **B)** Generalisation latency across the 10 successes in which significant improvement in solving latency was detected across the first 5 solving latencies. **C)** the first memory (recall) latency compared with the first and very last original latency (seconds) (N = 14) as well as the first and last generalisation latency (seconds) (N = 13). **D)** Memory latency across the first 5 successes. *P < 0.05.

When the problem solvers solved the generalisation task repeatedly, they decreased their generalisation latency across successes (GLMM: Z = −4.76, P <0.001, Fig. 4B). A significant improvement in the generalisation latency was first detected on the 3^rd^ success (1st vs. 3rd success: *Z* = −2.87, P = 0.004) and the generalisation latency to each success remained low from the 5^th^ success onward (Note S4). These results suggest that the problem solvers were still learning the solution before the 5^th^ success. Generalisation latency across successes was affected by the first solving latency in the original task (P = 0.002) and areas of green coverage (P = 0.008). Lower generalisation latency across successes (i.e., faster learning speed) was associated with lower first solving latency in the original task and less green area.

For the memory task, 20 (out of 38, 53%) innovators from 9 sites returned to the memory task. Of these 20 innovators, 14 (70%) of them solved the task successfully on their first or subsequent return, and thus were the ‘problem solvers’ for this task. The success of solving this memory task was affected by indirect human disturbance (P = 0.019), though it was also marginally affected by the proportion of success at the site level in the original task (P = 0.055) (Table 2a). When the problem solvers re-encountered the same task after an extended period of time, they used the same successful solution to solve this task. Their first memory latency (M = 6.8 ± 1.85s) was significantly lower than their first original latency of the original task (GLMM: *Z* = 2.27, P = 0.023) and comparable to their last original latency (*Z* = −1.01, P = 0.313, Fig. 4C) as well as their first and last generalisation latency (first: *Z* = 0.62, P = 0.535; Last: *Z* = −1.67, P = 0.095). These results suggest that they can quickly apply the learned successful solutions to solve the same problem.

The problem solvers’ first memory latency was significantly affected by their first and last solving latency in the original task (first: P = 0.020; last: P < 0.001), the last generalisation solving latency (P = 0.035) as well as the number of successes obtained in both of the original and the generalisation tasks (P = 0.012). Lower first memory latency was also predicted by increased indirect human disturbance (P = 0.002) and decreased green coverage (P = 0.050), though increased squirrel population size tended to be a factor (P = 0.051) (Table 2b). When the problem solvers repeatedly solved the same problem after an extended period of time, they did not show further improvement across successes (P = 0.149, Fig. 4D). The memory latency across successes was only predicted by the last solving latency in the original problem (P = 0.033) (Table 2c).

## Discussion

Few studies have already demonstrated urban environmental characteristics affect wildlife’s innovative problem-solving ability (Chow et al., 2021; Cook et al., 2017). Here, we proposed the ripple effect hypothesis that states that due to the shared underlying cognitive mechanisms, the initial effects that urban environment characteristics have induced on innovative problem-solving performance would continue to affect the performance of other related cognitive processes. We used a generalisation and a memory task that share underlying cognitive mechanisms (learning and memory) with the original task to test this hypothesis. We related the affected initial solving performances (of population and individual level) of the original task to the solving performances of the generalisation and memory tasks. Our results provide more detailed information to refine the hypothesis; the initial effects that urban environmental characteristics have induced on the innovative problem-solving performance continue to affect solving performance of related processes at individual level but not at population level. These affected performances include individuals’ generalisation latency across successes and their first memory latency. Urban environmental characteristics such as (direct and indirect) human disturbance affect the solving performance of related processes both at site and individual level. Together, these results suggest that the impacts of urban environmental characteristics on cognitive processes are far reaching; its initial impacts on a single cognitive ability can induce a more global impact on other related cognitive processes.

Costs are often associated with novel object exploration (e.g., Mettke-Hofmann et al., 2006). However, information that is gathered from exploring novel objects has been shown to facilitate solving novel problems (e.g., Christensen et al., 2021; Mettke-Hofmann & Gwinner, 2003), or allow urban wildlife to exploit novel resources especially for diet and habitat generalists (Ducatez et al., 2018) such as this species of squirrels (Krauze-Gryz & Gryz, 2015). In the case of solving novel problems, learning to solve a novel problem would presumably increase foraging efficiency if the learned information can be applied in a similar or the same context. Seemingly, the fact that the problem solvers show comparable first generalisation latency to their first original latency as well as showing signs of learning the solution for the generalisation task in the first few successes have suggested otherwise. However, the problem solvers became significantly faster to apply the learned solutions to solve the generalisation task on their third success. Moreso, the problem solvers were able to recall relevant information the first time when they saw the original task again as well as efficiently deploy the learned solution in the first and subsequent visits, resulting in the first memory latency being comparable to their last solving latency in the original task. In particular, more previous successful experiences in relevant tasks facilitates a problem solver to recall the learned successful solution faster when they encounter the same task. These results show the advantages of learning and memory in adapting to urban environments.

Problem-solvers’ first original latency has affected their generalisation latency across successes as well as the first memory latency. The first original latency was recorded as the time spent on manipulating a lever until the first success occurred. Crucially, this first original latency was measured as an accumulation of one or more failed attempts that was taken *before* a problem solver had learned the successful solution. Thus, the first original latency of the original task may entail other cognitive processes (e.g., attention) and non-cognitive traits (e.g., neophobia, motivation) that are not measured in this study but that have been shown to be related to problem-solving performance. For example, higher motivation or persistence, measured as the attempt rate, has been shown to increase efficiency in solving the same food-extraction apparatus in a closely related squirrel species, the Eastern grey squirrel, *Sciurus carolinensis* (Chow et al., 2016). These unmeasured cognitive processes and non-cognitive traits may reflect how the problem solvers react to their environment, and specifically to the environmental characteristics that we have measured. While we cannot pinpoint to which processes or non-cognitive traits is the first to be affected by the urban environmental characteristics in the original task, the first original latency appears to be the start of the ripple effect that continues to affect how the problem solvers receive feedback from their interactions with the task. This (positive) feedback during problem solving can influence their subsequent problem-solving performance (Cooke et al., 2021; Johnson-Ulrich et al., 2018).

Urban environmental characteristics have been previously shown to affect the innovative problem-solving performance of red squirrels, both at population and individual levels (Chow et al., 2021). As expected, these environmental characteristics also affect the generalisation and memory performance at both levels. Increased (direct and indirect) human disturbance decreases success rate of the generalisation task and the memory task at population level whereas decreased green coverage lowers the problem solvers’ generalisation latency across successes. Instead of highlighting which environmental characteristics are protective factors for urban wildlife (see Chow et al., 2021; Morand-Ferron & Quinn, 2011), our findings highlight that increased human disturbance and less area of green coverage are risk factors for urban wildlife. This suggestion can be supported by the innovators (who had solved the original task multiple times) residing in the site with the highest human disturbance (i.e., university) and the site with the least green coverage (i.e., Azusa park) did not participate in either the generalisation or the memory tasks. In these sites, it is likely that there is a higher risk of exposing themselves to potential threats (e.g., humans with dogs) even though the generalisation and memory tasks can merely be solved by using the learned solution. This result is striking as it suggests that some environmental characteristics like (direct and indirect) human disturbance pose a restriction on using the learned information. In other sites where the innovators have participated in the study, the enhanced learning and recall speed is likely a result of adaptive trade-off with the costs associated with these ‘harsh’ urban environmental characteristics (less green coverage, increased direct and indirect human disturbance). Because ‘disturbance’ entails many aspects (e.g., intensity, types of activity, distance toward humans) (Tablado & Jenni, 2017), more investigations are needed to reveal what aspects of human-related factors that wildlife reacts to, and their relationships to cognitive performance.

Overall, our results provide support for the proposed ripple effect hypothesis. We note that the natal dispersal of this species has limited the sample size for more detailed analyses (e.g., accumulated effects of each environmental characteristics, or interactions between urban environmental characteristics and affected performances in the original task on performances of the generalisation and memory tasks). Nevertheless, we have shown the direct effects of affected performance as well as identified relevant urban environmental characteristics that shape cognitive performance. Some of the urban environmental characteristics appear to induce a more global impact on cognitive processes. Future studies can examine similar questions and expand the investigations into the interaction effects of urban environmental characteristics (Isaksson, 2018) on cognitive performance. Ideally, investigations should be carried out in a species that has longer site fidelity. Such investigations will provide a more comprehensive picture when understanding the role of urban environmental characteristics in relation to cognition as well as gain insights into the mechanisms that support wildlife species adapting to urban environments.

## Supporting information

Supplementary materials that include additional analyses, tables, notes and videos

## Author Contributions

PKYC conceived, designed and carried out the experiment, ran the data analyses and wrote the first draft of the manuscript, KU and IK contributed significantly to the project.

## Open data access

All raw data can be downloaded from OSF ([here](https://osf.io/ufbvx/?view_only=af34ecb913f94d348e28f69e869295cf)).

## Acknowledgements

We thank Obihiro City and H. Yanagawa at Obihiro University for granting the study areas, O. Ryunosuke for logistic support, T Temizyürek for comments.

