## Supplementary materials that include additional analyses, tables, notes and videos for "‘Ripple effects’ of urban environmental characteristics on cognitive performances in Eurasian red squirrels"

**This PDF file includes:**

Supplementary Notes S1 and S4
Table S1 to S10
Legends for Videos S1 and S2
SI References

**Other supplementary materials for this manuscript include the following:**
Videos S1 and S2
Datasets S1 to S5

**Note S1. Individual identification and the number of squirrels in each site**

To identify each individual as well as record the number of squirrels in each site, we used an established method (1) alongside mark-recapture and mark-resight method. We subjected each video footage to a frame-by-frame analysis using Adobe Premiere Pro CS6. These video footage were records of video cameras that were set 1 m away from the apparatus in each field site. When we saw a squirrel the first time, it was 'marked' using their individual characteristics. It was 'recaptured' when it reappeared in the subsequent videos. We recorded each squirrel's characteristics as detail as possible that included their facial marking (e.g. a white dot/patch on face), colouration (e.g. orange, burgundy, brown patch on forehead), colour of limbs (e.g. orange/dark brown paws or dots on a toe) alongside height (relative to the apparatus), tail and body shape (e.g. full fur tail, half tail).

The first identification required intensive observer training that lasted for two months. Each squirrel was assigned a name and an identification number. This process required back and forth watching different footage so that the individuals' full characteristics could be revealed from different angles. We reidentified the squirrels three to five months after the first identification during which the same coder re-conducted the frame-by-frame analyses of the unmarked individuals by randomly selecting 3 out of 6 sites (total 23 individuals). To examine agreement between the two measures, we ran an intra-rater reliability test using Cohen's Kappa (Kappa = 0.99).

For marked individuals, we cross-checked the identity of individuals (with ear-tagged and collar) in five (out of 11) study sites that had ongoing field survey and trapping records (Table S1). Mark-resight method was used to confirm our observation. This method required walking slowly on the pathway in a site, or standing still in bushes or among trees to observe the squirrels. A squirrel was 'marked' using its unique characteristics mentioned above. We noted 'resight' when we saw the same squirrel. The double methods used here allowed us to check the identity of individuals in each site as well as to include the squirrels that were on trees.

**Note S2. Detailed measurements of each urban environmental characteristic.**
Chow and colleagues (2) have identified four urban environmental characteristics that were important for squirrels. These included the number of humans in a site (direct human disturbance), number of buildings around a site (indirect human disturbance), green coverage, and squirrel population size.

*Direct human disturbance.* We recorded the number of humans in a site 4-5 times daily regardless of weather conditions. In each record, we noted the time, weather, and the number of humans in a site before we set up the experiment in a site, before or after re-bating the apparatus, and when the experiment ended for the day, and thus resulting in 4-5 data points per day. Each record was obtained either from in-site recording or using distance-based methods. In-site recordings included walking around the site and counting each human that walked past the experimenter. This allowed us to count the number of humans more accurately if a site had a large area of bushes. Distance-based records meant dividing a site into four roughly equal areas and counting the number of humans in each area at the centre of the site or at the outer edge of the site, which minimises double counting. As there were other behavioural experiments that continued after the squirrels participated in the novel problem, we continued to obtain data on the number of humans in each site daily. We carried out an average of 38 observation days in each site (ranging from 33-42 days). We divided the total number of humans across the daily 4-5 scans in a site and across all observation days by the number of observation days to obtain the mean number of humans in a site per day.

*Indirect human disturbance.* Indirect human disturbance was the the number of human-built structures (e.g. schools, houses, restaurants and stores) within and 50 m surrounding each site. This 50 m covered urban red squirrels' minimum routine movement (3) where they can encounter anthropogenic food as well as capture the greatest human-induced disturbance as in different pollutants (e.g. noise, traffic, household and other human activities).

*Green coverage.* Green coverage of each site was defined as the areas covered by trees ($m^2$); a major resource that provides safety and food for squirrels. By using satellite mode of Google Map, we used point-to-point method, adding points around an area's boundary on the map and calculate the areas that were covered by trees site size ($m^2$)

*Squirrel population size.* Squirrels population size of each site was obtained using mark-recapture and mark-resight methods (see Note S1) that allowed us to double check the number of squirrels that we saw in the video footage and in each site.

**Note S3.** Detailed experimental procedures.

The field experimental protocol followed Chow and colleagues (2). Between May 2018 and January 2019, we tested 38 urban red squirrels in 11 sites (> 800 m between sites to avoid pseudo-replication) at Obihiro city, Hokkaido, Japan (see table S1 for site information). These sites were located at different places of the city, and vary with environmental characteristics of interest (e.g., direct and indirect human disturbance, areas of green coverage, squirrel population size). In these sites, squirrels are predominantly fed on Korean pine (*Pinus koraiensis*) and Manchurian walnut (*Juglans mandshurica* var. *sachalinensis*) trees.

The 38 squirrels were innovators who had previously repeatedly solved a novel food-extraction problem (i.e., the original problem, Fig. 1B). All squirrels were identified by an established method (1) that required frame-by-frame analysis of their characteristics from video footage as well as mark-recaptured and mark-resight methods (see Note S1). In addition to this, in five sites, the squirrels could be identified by their ear tags and/or collars from our long-term behaviour and population monitoring (4, 5).

3-5 days before starting the main experiment, we used hazelnut kernels to attract the squirrels to a location that was far away from major roads as well as close to a tree; this aimed to minimise road kill or predation risk (6). Once squirrels visited the location regularly (indicated by direct observation and emptied hazelnut kernels), we then presented the food-extraction problems (one box at a time) to the squirrels daily, during their most active period (from dawn to noon) to minimise possible confounding variables such as season, weather or motivation on performance.

During the experiment, we set the apparatus at where the original problem was and checked (i.e., rebait) it 3-4 times per day (45 mins-1.5 hours between checks). This inter-trial interval allowed us to to minimise social interference (less than 1% video had more than 1 squirrel on the apparatus at the same time) as well as recruit subordinate individuals when dominant individuals were at rest, and thus increased participation rate. In each check, we randomised the facing direction of the nut containers and the apparatus. For the generalisation task, the presentation sequence of the coloured levers was also randomised. For the memory test, which levers had a nut was chosen at random.

The field experiment lasted around 3.5 weeks in each site. All the innovators received the generalisation task the next day after they had completed the original problem. Before we presented the original problem to the innovators again, we carried out other behavioural assays that did not involve any similar solutions for solving the generalisation problem or the memory test. 21 days after the generalisation test, we presented the original problem to the innovators only for one day (from dawn to noon during their most active time); note that this has resulted in fewer successes obtained by each innovator in the memory test than in the generalisation problem.

**Note S4. Detailed results of the generalisation latency across successes**

In the generalisation problem, the repeated innovators showed significant improvement across the generalisation latency (GLMM $Z$ = -4.76, P < 0.001). Accordingly, we further examined the innovators' latency by conducting success-by-success comparisons. We found that compared with their generalisation latency on the first success (i.e., their first generalisation latency), the repeated innovators showed their first significant improvement on the 3$^{rd}$ success (1$^{st}$ vs. 3$^{rd}$: $Z$ = -2.87, P = 0.004) and they consistently solved the problem using low latency from the 5$^{th}$ success onward (1$^{st}$ vs. 5$^{th}$: $Z$ = -2.66, P = 0.008;1$^{st}$ vs. 6$^{th}$: $Z$ = -3.25, P = 0.001;1$^{st}$ vs. 7$^{th}$: $Z$ = -2.38, P = 0.017;1$^{st}$ vs. 8$^{th}$: $Z$ = -2.94, P = 0.003; 1$^{st}$ vs. 9$^{th}$: $Z$ = -2.76, P = 0.006; 1$^{st}$ vs. 10$^{th}$: $Z$ = -3.86, P < 0.001)

**Table S1.** Detailed information for the 11 study sites. This table is taken from Chow and colleagues (2021). Information include location (site name), using satellite mode of Google Map for GPS coordination and site size ($m^2$), surface area covered by tree canopy divided by site size (proportion of green area), participation rate in the original problem (the number of squirrels participating in the study divided by the actual squirrel population size in each site using mark-recapture and mark-resight methods) of each study location, and the number (and the type) of potential non-human predators recorded upon spotting one.

| Location | GPS coordination | Site size ($m^2$) | Green area (proportion) | Squirrel population size (Participation rate%) | Potential non-human predators (e.g., raptors, foxes, domestic cats and dogs) |
|---|---|---|---|---|---|
| Manabino park | 42.87, 143.19 | 46,433.4 | 0.34 | 9 (88.9%) | 3 (foxes, cats, raptors) |
| Riverside | 42.88, 143.18 | 68,162.6 | 0.71 | 9 (88.9%) | 3 (foxes, raptors, cats) |
| Azusa park | 42.93, 143.17 | 20,957.5 | 0.33 | 4 (100%) | 2 (cats) |
| Nishiobihiro park | 42.91, 143.13 | 40,686.1 | 0.60 | 9 (100%) | 3 (foxes, cats and dogs) |
| Tsuda park | 42.92, 143.12 | 85,769.6 | 0.74 | 7 (100%) | 3 (cats and dogs) |
| Fushikobetsu park | 42.92, 143.13 | 49,572.16 | 0.80 | 8 (87.5%) | 2 (cats and dogs) |
| Ishio Ryokuchi park | 42.91, 143.15 | 24,262.6 | 1 | 9 (100%) | 4 (cats) |
| Oyama Ryokuchi park | 42.90, 143.17 | 51,183.6 | 0.85 | 9 (100%) | 3 (cats) |
| Obihiro Forest (Baseball field) | 42.88, 143.15 | 171,859.5 | 0.73 | 6 (83.3%) | 2 (foxes, cats) |
| Obihiro University | 42.87, 143.17 | 388,390.6 | 0.09 | 5 (100%) | 2 (cats) |
| Ozora park | 42.88, 143.15 | 51,012.2 | 0.56 | 4 (50%) | 2 (cats and dogs) |

Legends for Videos S1 and S2.

**S1.** A squirrel, Mario, was solving the original problem (see here), which was also used for the memory test.

**S2.** The squirrel, Mario, was solving the generalisation problem (see here), a task that was similar but novel to the original problem. We changed the shape of the apparatus and the colour of the levers to maximise the differences between the original problem and the generalisation problem. This task was used to assess the innovators' ability to apply a learned solution of the original problem to solve the generalisation problem.

Legends for Datasets S1 to S5.

S1. Dataset about participation and success rate at site level (S1. Success_Participation_datasets)

S2. Dataset of the first and last innovation latency as well as the first generalisation latency (S2. Generalisation_Paired_latency)

S3. Dataset of generalisation latency across successes (S3. G_first and across_solving latency)

S4. Dataset of the first and last innovation latency as well as the first memory latency (S4. Memory_paired_latency)

S5. Dataset of memory latency across successes (S5. M_first and across_solving latency)
